# dRFEtools: Dynamic recursive feature elimination for omics

**DOI:** 10.1101/2022.07.27.501227

**Authors:** Kynon JM Benjamin, Tarun Katipalli, Apuã CM Paquola

## Abstract

Technology advances have generated larger omics datasets with applications for machine learning. Even so, in many datasets, the number of measured features greatly exceeds the number of observations or experimental samples. Dynamic recursive feature elimination (RFE) provides a flexible feature elimination framework to tackle this problem and to gain biological insight by selecting feature sets that are relevant for prediction. Here, we developed dRFEtools that implements dynamic RFE, and show that it reduces computational time with high accuracy compared to RFE. Given a prediction task on a dataset, dRFEtools identifies a minimal, non-redundant, set of features and a functionally redundant set of features leading to higher prediction accuracy compared to RFE. We demonstrate dRFEtools’ ability to identify biologically relevant information from genomic data using RNA-Seq and genotype data from the BrainSeq Consortium. dRFEtools provides an interpretable and flexible tool to gain biological insights from omics data using machine learning.

## Background

The power of next generation high-throughput sequencing technologies is unquestionable. The creation of increasingly larger epigenetics, genetics, and transcriptomic datasets has generated more comprehensive insights into human biology. The potential of omics (transcriptomics, genomics, methylomics, etc.) expands from discovery biology to identification of gene targets for the development of biomarkers and novel therapeutics for various diseases as reviewed in[1, 2]. Even so, these technological costs and sample availability generates increasingly more features compared to samples. This large *p* small *n* problem can result in overfitting[3]. As such, feature selection is essential to solving this problem.

Recursive feature elimination (RFE) is one such feature selection method first proposed in conjunction with SVM for binary class problems[4], which has since been extended to multi-classes[5, 6] and additional algorithms[7, 8]. This iterative process is an instance of backward feature elimination, which optimally removes one feature at a time. For computational considerations, we can eliminate a substantial number of features (feature subset ranking); however, it can be difficult to balance computational time (small number of features dropped) and model performance degradation (substantial number of features dropped). To overcome this issue, RFE can be done dynamically, which provides a more flexible feature elimination operation by removing a substantial number of features at the beginning and becoming a single feature elimination when there are a small number of features. While the framework for dynamic RFE has been discussed[9, 10], to the best of our knowledge there is no software available that implements dynamic RFE.

Here, we developed a python package for dynamic RFE, dRFEtools, and applied it on simulated data and three biological scenarios using a subset of the postmortem dorsolateral prefrontal cortex (DLPFC; n=521) from the BrainSeq Consortium[11, 12]. In addition to providing helpful functions to optimize, implement, and plot dynamic RFE, dRFEtools decreases computational time compared to the current RFE function available with scikit-learn, a popular platform for machine learning in python, while maintaining high accuracy in simulated data for both classification and regression models. Furthermore, we demonstrate the utility of dRFEtools as an orthogonal approach to commonly used bioinformatics methods with an additional ability to prioritize a smaller subset of genomic features that contain biologically relevant information.

## Results

### dRFEtools: dynamic recursive feature elimination

#### Framework for scikit-learn integration

The purpose of dRFEtools is to implement dynamic RFE with scikit-learn for classification and regression using linear models, random forest, and support vector machines (SVMs). As feature selection is essential for omics data, where there are far more features compared to samples, dRFEtools seeks to provide a utility that is fast and accurate for big omics feature selection. The development of dynamic recursive feature elimination (**Figure 1A**) for scikit-learn models can be separated into two main parts: 1) feature iterator (**Figure 1B**), and 2) model evaluation for *N* features (**Figure 1C**). For the feature iterator (**Figure 1B**), we use *keep_rate* (*1 - elimination_rate*) to determine the percentage of features to drop for each iterative operation. This outputs the number of features (*nf*) that should remain after each iterative operation.

**Figure 1:**
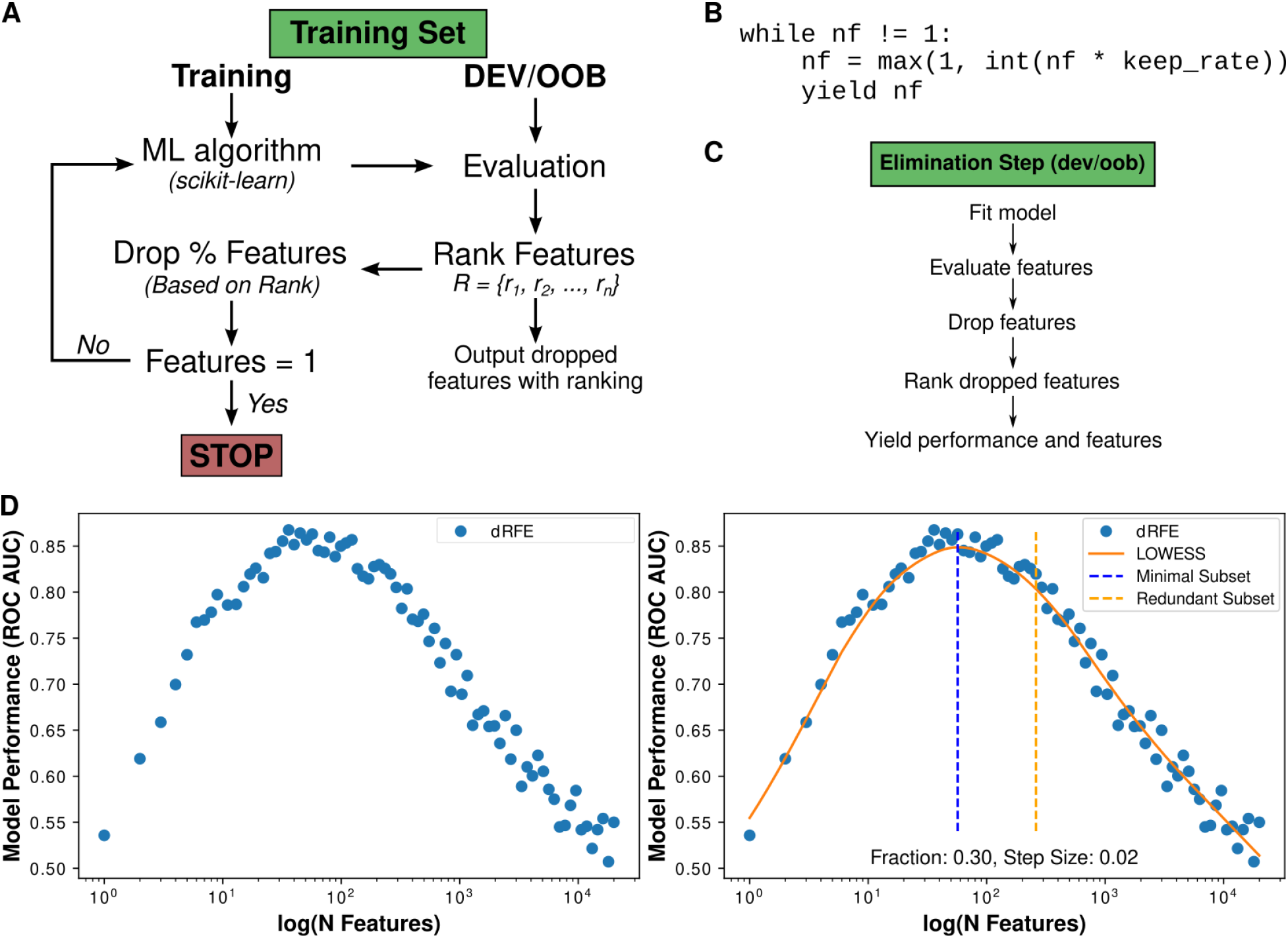
Schematic of dynamic recursive feature elimination for dRFEtools. **A.** Flowchart showing recursive elimination process, where scikit-learn model can be either classification or regression. We ranked the dropped features and saved them for downstream analysis. **B.** Feature iterator code used to generate dynamic elimination. **C.** Flowchart showing the elimination steps using the developmental or OOB set. **D.** Example of LOWESS fitting on dRFEtools model performance multiple classification using simulated data using area under the receiver operating characteristic curve (ROC AUC). Simulated data generated for multi-classification using scikit-learn *make_classification* for four classes (500 samples, 20,000 features, 100 informative, and 300 redundant). Solid orange line - LOWESS fit. Dotted orange line - estimated redundant subset of predictive features. Dotted blue line - estimated smallest subset of predictive features. Blue dots - results from dRFEtools.

The Elimination Step process (**Figure 1C**) can be broken down into five steps. In the first step, we fit our classification or regression model to the training set. The features are then evaluated based on a feature importance score estimated from the trained model. For linear models and SVM, the absolute weights (|w|) of each feature are used directly. For random forest, the unsigned feature importance (Gini) is used to evaluate features. This improves computational time and performance with correlated features compared to permutation importance[13]. Following this evaluation, we select the features with the lowest importance to be dropped based on the elimination rate. We rank and output these low importance features by default. In the last step, we use the Out-of-Bag (OOB) for the random forest models or developmental set −20% of the training set - for all other algorithms to measure model performance. We return these measurements (*N* of features, model performance, remaining features) for each iteration. To reduce the problem of overfitting, we recommend using *n*-fold cross validation with this method.

To evaluate model performance during dynamic RFE, we measured three metrics for classification and regression. For classification analysis, we evaluated features at each step for normalized mutual information (NMI), accuracy, and area under the ROC (receiver operating characteristic) curve. dRFEtools automatically detects if the target is binary or multiple classes and adjusts parameters accordingly. For regression analysis, we evaluated features at each step for r^2^ correlation, mean square error (MSE), and explained variance.

#### Optimal feature selection for smallest and redundant subsets

Genetic redundancy, where two or more genes perform the same function, is widespread in the genomes of higher organisms [14]. As such, it is often biologically relevant to select a subset of features that are functionally redundant. To this end, dRFEtools outputs both the smallest and redundant subset of features that holds predictive power.

From the locally weighted scatterplot smoothing (LOWESS) curve (see **Methods**), we extract the local maximum as the smallest subset of features and return the number of features closest to this number (**Figure 1D**). To determine the redundant subset of highly predictive features, we examine the rate of change of the lowess curve and select the point at which the slope changes steeply. From this, we return the number of features closest to this number. An optimization function is available in dRFEtools to help determine the best parameters to select the smallest and redundant subset of features (**Data S1**).

### dRFEtools outperforms current RFE algorithm for feature selection

To assess the ability of dRFEtools to accurately identify informative features in classification and regression problems, we performed two different simulation analyses to compare dRFEtools with the current RFE scikit-learn function using four algorithms for classification (logistic regression, random forest classifier, stochastic gradient descent, and support vector classification) and regression (ridge, elastic net, random forest regressor, support vector regression) and measured: 1) feature selection accuracy, 2) feature selection false discovery rate (FDR), and 3) computational time.

For our first simulation analysis, we generated simulation classification and regression data using scikit-learn with two elimination rates for RFE (10% and 100 features) and dRFEtools (10% and 20%). For RFE, 10% represents a static number equal to 10% of total features (2000 features each iteration). For dRFEtools, the number of features eliminated each run was determined dynamically each iteration. Using these elimination rates, we found dRFEtools dynamic approach improved accuracy and decreased false discovery rate for both classification and regression (Two-way ANOVA, p-value < 0.01; **Figure 2A,B**) as well as higher area under the ROC curve for feature selection (**Figure S1**). Notably, the random forest classifier and regressor algorithm showed the lowest FDR as well as the highest feature selection accuracy for both RFE and dRFEtools. Even so, computational time and FDR were significantly reduced for the dRFEtools classification and regression models compared to RFE (One-way ANOVA, p-value < 0.01).

**Figure 2:**
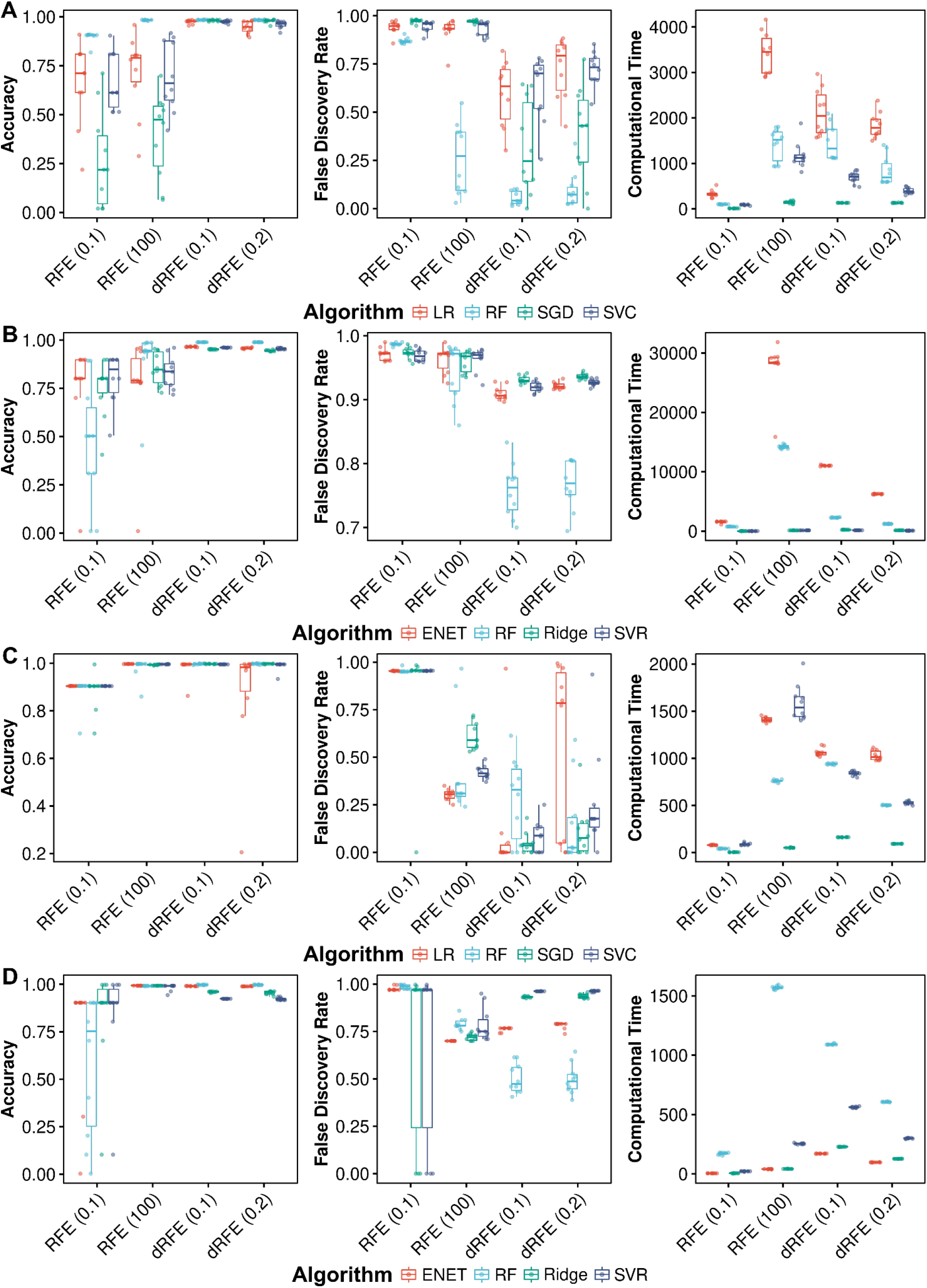
dRFEtools is fast and accurate at selecting informative features compared to current RFE algorithms. Simulations with 10-fold cross-validation of **A.** classification and **B.** regression with scikit-learn sample generators shows dRFEtools has high feature selection accuracy and low false discovery rate compared to standard RFE. Computational time is also decreased compared to more accurate RFE (100 step elimination). Biological **C.** classification and **D.** regression simulations (see **Methods**) with 10-fold cross-validation show similar high accuracy and lower false discovery rate of dRFEtools compared with standard RFE. For biological classification, we generated 10 simulated bulk RNA-sequencing dataset (20k genes, 400 samples, and 100 informative genes) with SPsimSeq [15] and raw counts from the Cancer Genome Atlas (TCGA) lung adenocarcinoma [16, 17]. For regression, we generated 10 simulated quantitative trait loci datasets (10k variants with an allele frequency between 0.05 and 0.95, 500 samples, and 30 informative variants) with PLINK [18, 19]. LR: logistic regression, RF: random forest, SGD: stochastic gradient descent, ENET: elastic net, SVC: support vector classifier, and SVR: support vector regressor. RFE (0.1): RFE with 10% representing a static number equal to 10% of total features (2000 features each iteration). RFE (100): RFE that eliminates 100 features each iteration. Accuracy: feature selection accuracy (**Equation 2**). FDR: feature selection false discovery rate (**Equation 3**). Computational time in seconds.

Next, we generated biologically relevant simulation classification (bulk RNA-sequencing) and regression (quantitative trait) data for our second simulation analysis. Again, we used the same elimination rates and machine learning algorithm as the scikit-learn simulations. Similar to the scikit-learn simulations, we found dRFEtools outperformed RFE for accurate feature selection (**Figure 2C,D** and **Figure S2**). Specifically, we found a decrease in FDR for all algorithms in classification and for random forest in regression (Two-way ANOVA, p-value < 0.05). Additionally, we found feature selection accuracy improved over the 10% RFE model similar to the scikit-learn simulation analysis. In general, random forest algorithms outperformed the other algorithms for accuracy feature selection (One-way ANOVA, p-value < 0.01).

For the quantitative trait simulation data, we simulated variants associated with a continuous phenotype similar to gene expression. As such, we also considered one-hot encoding for our simulated variants as features. This improved accuracy for the Ridge and SVR algorithms using dRFEtools at both elimination rates (**Figure S3**). The significant increase in features greatly affected computational time for the RFE 100 step elimination, but not dRFEtools. Additionally, FDR decreased with RFE for the elastic net algorithm. Altogether, suggesting that using one-hot encoding for variants is application and algorithm dependent.

### Applications of dRFEtools to biological data

To illustrate dRFEtools application to biological data, we considered a subset of data from the BrainSeq Consortium Phase 1 DLPFC adult (age > 17) postmortem brain collection (n=521) [11]. With this dataset, we considered three scenarios: 1) binary classification for schizophrenia (n=172) and major depression disorder (MDD; n=142) using gene expression from poly-adenylated RNA-sequencing as features, 2) multi-class classification of neuropsychiatric disorders (neurotypical control, n = 207) using gene expression as features, and 3) regression modeling to impute gene expression using SNP genotypes as features.

#### Binary classification of neuropsychiatric disorders

For binary classification for schizophrenia and MDD, we generated two subgroups including neuropsychiatric disorder of interest (schizophrenia or MDD) and neurotypical controls. With these two subgroups, we used dRFEtools to apply random forest classification to predict diagnosis compared with control with 10-fold cross-validation. Here, we found 45 features showed a high test prediction accuracy for MDD (median of 82.8%) compared with the 56 features that were predictive for schizophrenia (median 68.4%) although there were 30 more schizophrenia samples (**Figure 3A** and **Data S2**). This was robust when we expanded the features to our “redundant” subset (Median test accuracy, 81.4% and 71.6% for MDD and schizophrenia, respectively; **Figure 3A**).

**Figure 3:**
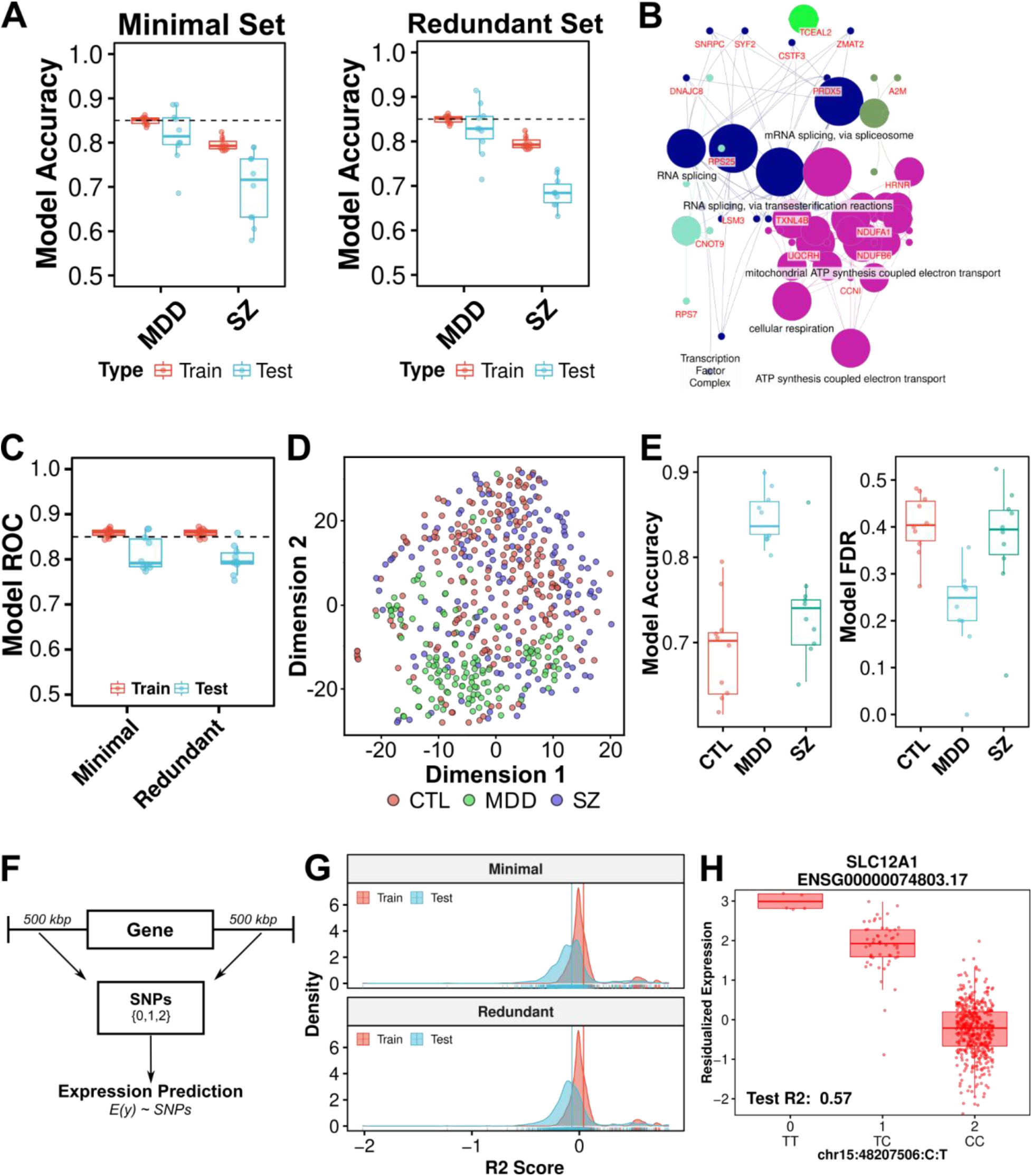
Biologically relevant features selected using dRFEtools associated with prediction accuracy. Binary classification of major depression disorder (MDD) and schizophrenia (SZ) compared with neurotypical controls (CTL) shows **A.** robust model accuracy for minimal and redundant feature sets as well as **B.** biologically relevant information. Multi-classification of neuropsychiatric disorder shows high **C.** model accuracy (ROC) driven by high classification accuracy and low false discovery rate (FDR) of MDD compared to CTL or SZ. **D.** tSNE plot demonstrating predictive features’ ability to separate MDD from CTL or SZ. **E.** Boxplot of model accuracy and FDR for correct diagnosis classification (one-vs-rest). Gene expression imputation from *cis*-SNPs using random forest regression selects genes with strong genetic variance association. **F.** Schematic showing expression imputation approach. **G.** Distribution of test r^2^ for all genes across folds for minimal and redundant SNP sets. **H.** Example of high median test r^2^ gene-SNP pair identified from dRFEtools random forest regression. 10-fold cross-validation used for all scenarios.

To understand the biological relevance of these features, we first examined the overlap of the smallest subset of predictive features with those extracted from traditional differential expression (DE). Of the more than 2000 genes dysregulated for MDD identified from DE analysis, 26 of the 45 (57.8%) were also among the smallest subset of predictive features (**Figure S4A**). In contrast, only 14 of the 56 genes (25%) among the smallest subset of predictive features were also among the 152 DE genes dysregulated for schizophrenia (**Figure S4B**). Interestingly, we found significant enrichment of MDD and schizophrenia in the smallest subset of predictive and redundant features compared to the DE analysis (Fisher’s exact test, p-value < 0.001), demonstrating the ability of dRFEtools to select biological relevant features even with low prediction accuracy. Even so, the prediction test score accuracy appeared to correlate with degree of biological relevance for features selected. Indeed, we observed significant gene term enrichment for mitochondria related terms with the redundant MDD predictive gene set (**Figure 3B** and **Data S3**), which is extensively associated with MDD [20].

#### Multi-classification of neuropsychiatric disorders

We next applied dRFEtools for a multi-classification scenario. To this end, we applied dRFEtools with random forest classifier and 10-fold cross-validation to classify diagnosis status (neurotypical controls, MDD, or schizophrenia). Here, we found a minimum of 71 features showed a high area under the ROC curve test prediction score for diagnosis classification (median of 81.3%) (**Figure 3C** and **Data S4**). Our redundant subset (n = 583 features) also showed a high ROC test prediction score (median of 80.0%; **Figure 3C**). Given the low prediction accuracy for schizophrenia binary classification, we examined the ability of the minimum predictive set (n=71) to cluster individuals based on diagnosis. Unsurprisingly, we found clear separation of MDD from neurotypical controls and schizophrenia, but little to no separation of schizophrenia from controls (**Figure 3D**). Indeed, when we examined the accuracy for predicting a specific diagnosis, we found a low accuracy and high FDR for predicting schizophrenia (Accuracy = 74.0% and FDR = 39.4%) and neurotypical controls (Accuracy = 70.2% and FDR = 40.4%) compared with MDD individuals (Accuracy = 83.7% and FDR = 24.9% of case/control prediction; **Figure 3E**).

#### Expression imputation regression model using genetic variants

Our third scenario applied dRFEtools in a random forest regression model with 10-fold cross-validation to select a small subset of single nucleotide polymorphisms (SNPs) that showed strong association with gene expression (i.e. imputing gene expression). To this end, we selected SNPs within 500 kbp up- and downstream the gene body of neuropsychiatric predictive genes (69 non-mitochondrial genes) from the second scenario (**Figure 3F**). While most genes did not show an association between gene expression and genetic variants (SNPs; **Figure 3G**), we found the six genes (*SLC12A1, PSPHP1, SH3PXD2A-AS1, TBC1D3L,* ENSG00000272977, and ENSG00000263667) with high association between expression and SNPs (median test r^2^ > 0.1; **Data S5**).

Similar to eQTL analysis, the six genes identified had at least some of their expression phenotype explained by genetic variance (**Figure 3H** and **Figure S5**), as such we next examined the individual SNPs genetic variance contribution to expression. To this end, we calculated the proportion of variation explained for each predictive SNP of gene expression per gene and found a single SNP explained between 13 and 66% (**Table 1**) with all six genes showing a significant correlation (Linear regression, Bonferroni p-value < 0.05) between SNP rank from dynamic RFE and variance explained. As we did not remove SNPs in high linkage disequilibrium (LD) prior to applying dRFEtools, we next examined LD of predictive SNPs for each gene. Although the majority of SNPs (>72%) for five of the six genes were not in LD (r^2^ > 0.6; **Table 1**), several variants that explained a large proportion of variances were in high LD (r^2^ > 0.9; **Figure S6**). For example, three SNPs accounted for the majority of variance explained for *SLC12A1* all of which were in high LD (r^2^ > 0.97).

**Table 1:** Summary of multi-classification predictive features with high correlation (median test r^2^ > 0) between genotypes and gene expression determined using dynamic RFE performed with dRFEtools and random forest regression. Median R2 - median test r^2^ from dRFEtools random forest regression). Percentage of maximum variance explained from a single SNP on gene expression. Total number of SNP predictive for gene expression as well as the percentage of SNPs in LD (r^2^ > 0.6) of each other within these predictive SNPs in parentheses. * Denotes predictive features that have significant correlation positive (Linear regression, Bonferroni < 0.05) between rank as determined by dRFEtools and variance explained for specific SNP.

| Features | Median Test R2 (%) | Variance Explained (single SNP; %) | Total SNPs (% in LD) |
| --- | --- | --- | --- |
| PSPHP1 | 63.3 | 39.2 | 105 (16.2) |
| SLC12A1 | 57.2 | 66.3 | 45 (8.9) |
| SH3PXD2A-AS1* | 48.3 | 51.2 | 58 (27.6) |
| ENSG00000272977* | 44.4 | 15.0 | 45 (44.4) |
| TBC1D3L* | 42.5 | 27.6 | 44 (6.8) |
| ENSG00000263667* | 10.6 | 13.0 | 83 (6.0) |

## Discussion

The increasing omics (transcriptomics, genomics, methylomics, etc.) datasets have allowed for more machine learning applications to answer biological questions or identify biomarkers or new therapeutics. Dynamic recursive feature elimination provides a flexible feature elimination framework for genomic large *p* small *n* problems. Here, we developed the python package dRFEtools that integrates with scikit-learn to implement dynamic RFE. We show that it reduces computational time with high accuracy compared to RFE currently implemented in scikit-learn in simulated data. Furthermore, we demonstrate its application to identify biologically relevant information from genomic data using the BrainSeq Consortium DLPFC adult (age > 17) postmortem brain collection (Phase 1; n= 521,[11]) on three scenarios: 1) binary classification for major depression disorder (n=142) and schizophrenia (n=172) compared with neurotypical control (n=207), 2) multi-classification of neuropsychiatric disorders, and 3) expression imputation using regression modeling. To the best of our knowledge, this is the first software implementing dynamic RFE.

We built dRFEtools around interpretability. Specifically, we implemented two key aspects to provide model transparency and interpretation: 1) automatic (default) generation of all features ranked and 2) LOWESS smoothing to extract both the smallest and redundant subset of predictive features. For optimization of the selection process, we provide several convenient functions that allow for visualization of the fitted curve as well as the ability for users to include more or less features within the redundant subset. In addition to this, we have also included support for numerous algorithms available in scikit-learn (e.g., linear models, random forest, and SVM). Thus, allowing for dRFEtools to be applied on binary classification, multi-class classification, and regression problems similar to RFE currently available in scikit-learn in contrast with other feature selection frameworks that are limited to classification problems[8–10, 21, 22].

While there is no current software available that implement dynamic RFE to compare dRFEtools with, we demonstrated dRFEtools ability to reduce computational time while maintaining or outperforming feature selection compared with scikit-learn implemented RFE. We found these results were robust using two kinds of simulated data: 1) scikit-learn simulated data and 2) biologically relevant (e.g., bulk RNA-seq or SNP-trait association). Of note, we found random forest classification and regression consistently outperformed other algorithms with false discovery rates significantly lower than other algorithms. More importantly, we found computational times were significantly decreased with regression problems showing the greatest reduction. This is essential as lower computational times allows for the expansion of expression correlation feature selection analysis to the genome-wide level with high accuracy, which is currently computationally prohibitive with traditional RFE.

In addition to simulated data, we illustrated dRFEtools applicability to neuropsychiatric disorder transcriptomics and genomic data. Here, we found that results from dRFEtools provided biological relevant information based on prediction accuracy, showing high overlap with traditional methods as well as enrichment for gene terms that are related to disease. Furthermore, we demonstrated the ability of dRFEtools to select SNPs significantly associated with gene expression with limited linkage disequilibrium. Additionally, dRFEtools was able to rank SNPs according to its contribution to expression variation showing the power of dynamic RFE.

## Conclusions

In conclusion, we have developed dRFEtools - a software that implements dynamic recursive feature elimination - with flexibility and interpretability in mind. We have shown its ability to substantially reduce computational time while maintaining high accuracy and low false discovery rate for feature selection in simulated data. Additionally, we have demonstrated dRFEtools ability to select features that contain biological relevant information in both classification and regression problems. With this software, the power of dynamic RFE is now more widely available to increase our ability to gain insights from omics data using machine learning.

## Methods

### Dynamic recursive feature elimination (dRFEtools) framework

#### Optimization of feature selection line fitting

For the optimization of feature selection for the smallest and redundant subsets, we first fitted a line to the data. For line fitting, we first interpolated the data by finding the B spline representation of log10 transformed features (x) and model performance (NMI for binary classification, ROC curve for multiple class classification, and r^2^ for regression) using *interpolate.splrep* from the SciPy library [23]. Given the set of data, a smooth spline approximation of degree *k* on the interval x_b_ <= x <= x_e_, where the degree of the spline is set to 3, x_b_ = x[0] and x_e_ = x[-1]. We then used the B spline to interpolate 5000 points between x_b_ and x_e_ to improve the quality of the fitted line. We used the LOWESS to create a smooth line of log10 features, *x*, against model performance, *y*.

#### Extracting redundant features

To determine the cutoff point for the number of redundant features based on the LOWESS curve, the rate of change was calculated using NumPy [24] as described in **Equation 1**, where *x* is the feature rank and *y* is the LOWESS smoothed curve. The rate of change between two points, *dxdy*[*i* + 1] – *dxdy*[*i*], was compared with the *step_size* parameter. We have set the default *step size* parameter at 0.02, but we suggest this should be optimized for each experiment. If the rate of change is positive and larger than the parameter set *step_size,* we annotated this feature rank. Once all features have been reviewed for a positive large rate of change (i.e., larger than step size), we select the highest ranking feature. As these are interpolated log10 estimated feature ranks, we find the model feature rank closest to this number and return the log10 transformed and original feature rank (i.e., untransformed).

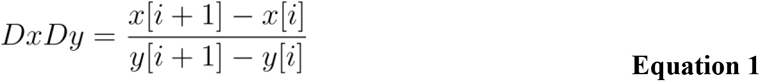

### Simulated data and analysis

#### Optimization and scikit-learn simulation analysis

For optimization examples and initial dRFEtools comparison with recursive feature elimination, we simulated classification and regression data using scikit-learn with the *make_classification* and *make_regression* functions. For binary classification, we generated data similar to high-throughput sequencing with 500 samples, 20,000 features, 100 informative and 300 redundant with one cluster per class for two classes. For multi-class classification, we generated data similar to binary classification modified for four classes (500 samples, 20,000 features, 100 informative, and 300 redundant). For regression, we generated data for one target using the same number of samples and total features as classification (500 samples, 20,000 features). Here, we increased informative features to 200 and introduced 20% bias and set noise to 0.5. For optimization bias was reduced to 2%.

#### Biological simulation analysis

For biological comparison of dRFEtools with recursive feature elimination, we simulated bulk RNA-sequencing data and quantitative traits loci from genotypes. For bulk RNA-sequencing simulation data, we used semi-parametric simulations with SPsimSeq [15] to generate 10 bulk RNA-sequencing data. To train the model, we downloaded raw counts of the Cancer Genome Atlas (TCGA) lung adenocarcinoma [16, 17] data using recount3 [25] and constructed an edgeR DGE object [26, 27]. As recommended by SPsimSeq, we filtered out lowly expressed counts. For this, we used *filterByExpr,* which keeps genes that have count-per-million (CPM) above 10 CPM in 70% of the normal tissue (smallest group). Once filtered, we used the *SPsimSeq* function to generate 10 simulated datasets with 20k genes, 400 samples (240 tumor, 160 normal), 100 true differentially expressed genes with a minimal log fold change of 0.10 in a single batch.

For quantitative trait loci simulation, we used PLINK (v1.90p) [18, 19] to generate simplistic variant simulation, specifically all variants simulated are unlinked and in linkage equilibrium. To this end, we used the *simulate-qt* command to simulate a quantitative trait based on 30 informative variants out of a total of 10k variants for 500 samples with an allele frequency between 0.05 and 0.95 for the trait-increasing allele. The effect size was generated to give a population variance explained of 0.03. From these frequencies, we simulated 10 different populations. For evaluation of dRFEtools, we used variants with and without one-hot encoding.

#### Metrics evaluation

To evaluate dRFEtools with RFE, we examined three metrics: 1) feature selection accuracy (**Equation 2**), 2) feature selection false discovery rate (**Equation 3**), and 3) computational time in seconds. We examined these metrics across four models for classification and regression and 10 fold cross-validation. For classification, we used logistic regression with L2 penalty and max iteration of 1000, support vector classification (SVC) using max iterations of 10k, stochastic gradient descent with perceptron loss and L2 penalty, and random forest classifier with Out-of-Bag scoring and 100 *n* estimators. Cross-validation folds were generated with *StratifiedKFold*. For regression, we used ridge with default parameters, elastic net with alpha equal to 0.01, support vector regression (SVR) using max iteration of 10k, and random forest regression with Out-of-Bag scoring and 100 *n* estimators. Cross-validation folds were generated with *KFold.*

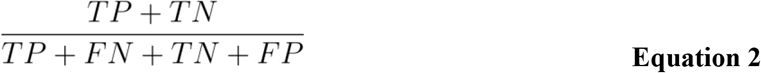

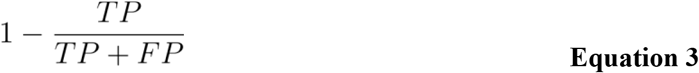

In addition to feature selection accuracy and false discovery rate, we also used the ROC curve (receiver operating characteristic curve) to assess simulation feature selection performance. To calculate the true positive (**Equation 4**) and false positive (**Equation 5**) rates for feature selection performance, we calculated the cumulative sum of true positives (TP) and false positives (FP) at each threshold, where the threshold was determined by the cumulative features. Features were ranked based on simulation ground truth.

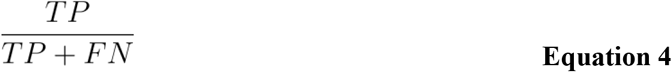

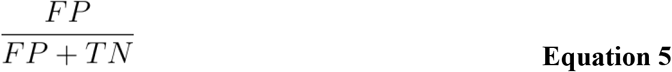

### Biological data and analysis: BrainSeq Consortium

For real data demonstration, we used the BrainSeq Consortium phase 1 gene expression and genotype data to model three biological scenarios: 1) major depression disorder (MDD) binary classification, 2) neuropsychiatric disorder multi-classification, and 3) expression imputation.

#### BrainSeq Consortium data availability

The original BrainSeq Consortium Phase 1 [11] processed gene level data mapped to hg19 can be downloaded as an *RangedSummarizedExperiment* R object from http://eqtl.brainseq.org/phase1/. This R variable contains raw counts, gene annotation (hg19) and phenotype data for neurotypical control and schizophrenia. The full data including MDD and bipolar disorder individuals were reprocessed to the hg38 annotation for comparison analysis in BrainSeq Phase 2 [28]. For this study, we used the reprocessed gene level BrainSeq Phase 1 data on hg38 built generated in BrainSeq Phase 2.

DNA genotype data can be downloaded through Globus [11]. Trans-Omics for Precision Medicine (TOPMed) [29–31] imputed genotypes using the were generated as part of BrainSeq Phase 3 [32] and includes a total of 11,365,460 common variants after quality control. Briefly, this quality control included removing variants with low imputation quality (R2 < 0.7), minor allele frequency (< 0.01), missing call frequencies (> 0.1), and Hardy-Weinberg equilibrium (< 1e-10) using PLINK2 (v2.00a3LM; [33]).

#### Sample selection

We selected samples using four inclusion criteria: 1) either African or European ancestry, 2) individuals age > 17, 3) diagnosis of neurotypical controls, MDD, or schizophrenia, and 4) TOPMed imputed genotypes available. This resulted in a total of 521 samples for the DLPFC. Subject details are summarized in **Table S1**.

#### Normalized and residualized expression

For each scenario, we generated normalized and residualized expressions as previously described [32]. Briefly, we constructed edgeR objects with raw counts and sample phenotype information from the *RangedSummarizedExperiment* R objects. Similar to the biological simulation, we filtered out low expression counts using filterByExpr from edgeR and applied voom normalization [34, 35] (**Equation 6**). For residualized expression, we regressed out covariates from voom normalized expression using a null model that did not include the variable of interest (**Equation 7**). Following regression, we applied z-score transformation. For binary classification, normalized and residualized expressions were generated in two subgroups containing neurotypical control and one of the neuropsychiatric disorders (MDD or schizophrenia). For imputing expression, residualized expression was generated using **Equation 6** to account for diagnosis expression differences.

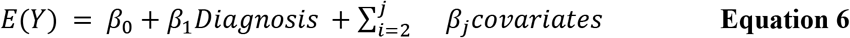

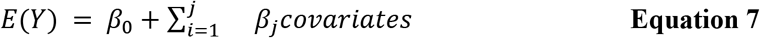

To determine covariates, we examined all non-boolean variables within the *RangedSummarizedExperiment* R object. From this, we removed variables that continued duplicate information (correlation > 0.95) using cytominer [36] function *correlation_threshold*. Following this, we selected variables that showed a correlation with expression before and after residualization. In place of testing all genes, we reduced dimensionality using principal components analysis (PCA) on voom normalized and residualized expressions. From this analysis, we selected sex, age, RIN, mitochondria mapping rate, rRNA mapping rate, genome mapping rate, adapter content, percent GC, population structure (genomic ancestry SNP PCs 1-3), and qSVs generated from BrainSeq Phase 1 [11, 37].

#### Differential expression analysis

To compare binary classification with traditional differential expression analysis, we fitted voom normalized counts using lmFit from limma (**Equation 6**) and identified differentially expressed genes using eBayes [38]. We performed this separately comparing neuropsychiatric disorder (MDD or schizophrenia) with neurotypical controls.

#### Dynamic RFE analysis

For all three scenarios, we used the random forest algorithm with Out-of-Bag scoring and 100 *n* estimators similar to simulation data. Additionally, we used *StratifiedKFold* to generate cross-validation folds for all scenarios to maintain even distribution of patient diagnosis across folds. We set the elimination rate to 10% with the default step size of 0.02 for all scenarios. For binary classification, we set 0.35 as the fraction of samples used for LOWESS smoothing, while multi-class and regression scenarios used the default of 0.3. For binary and multi-class classification, we performed on-the-fly residualization of expression to reduce potential leakage from training to test set in the residualization process. Specifically, we fit our null model to the training set and extracted coefficients to be regressed out of both training and test set expression.

#### Gene term enrichment analysis

For gene term enrichment analysis, we used GOATOOLS Python package [39] using hypergeometric tests with the Gene Ontology database. For network visualization, we used ClueGO (v2.5.8) [40] in Cytoscape (v3.9.1) [41].

#### Expression imputation preparation and linkage disequilibrium analysis

For expression imputation, we extracted SNPs that were 500 kbp up- and downstream the gene body using PLINK2. Residualized gene expression (**Equation 6**) was added to the SNPs data frame selected for a specific gene. We calculated linkage disequilibrium for genes with a median test r^2^ > 0.1 using PLINK for all predictive SNPs per gene.

#### Genetic contribution to expression

To examine the genetic contribution of the predictive SNPs to gene expression, we calculated the proportion of variation explained by adding one of the predictive SNPs to the linear model (**Equation 6**) using coefficient of partial determination through a relative variation index (**Equation 8**) as previously described [42]. Specifically, we used the residualized expression to remove the effects of all model covariates to generate reduced expression per gene. We then built the relative variation index (Δ*Pst*) for each gene comparing the reduced model (*Pst*, **Equation 9**) and the full model by including an individual SNP (*P_st_cis__*, **Equation 10**) from the predictive SNPs obtained from dRFEtools:

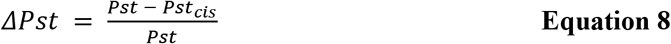

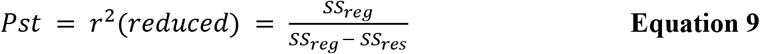

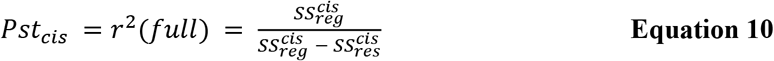

where *SS_reg_* denotes the regression sum of squares and *SS_res_* denotes the residual sum of squares.

### Graphics and code availability

The dRFEtools generates optimization plots using matplotlib [43, 44] in python and feature elimination plots with plotnine - a python implementation of ggplot2 [45]. We also generated tSNE plots in python with plotnine. All other boxplots and scatterplots were generated in R (v4.0.3) [46] using ggpubr (v0.4.0) [47]. Venn diagrams were generated with ggvenn [48].

dRFEtools is available on Python Package Index (PyPI) https://pypi.org/project/drfetools/ or on GitHub at https://github.com/paquolalab/dRFEtools. The code and jupyter notebooks that produce the results for this manuscript are available through GitHub at https://github.com/LieberInstitute/dRFEtools_manuscript.

## Supporting information

Supplementary Data 1: Optimization jupyter notebook

Supplementary Data 2: Binary classification

Supplementary Data 3: MDD GO analysis

Supplementary Data 4: Multiclass analysis

Supplementary Data 5: Gene imputation

## Declarations

### Ethics approval and consent to participate

Not applicable.

### Consent for publication

Not applicable.

### Availability of data and materials

The gene expression datasets analyzed in the current study aligned to the hg38 [28] are available upon request. The original hg19 aligned R variables with neurotypical control and schizophrenia individuals is available at http://eqtl.brainseq.org/phase1/. The FASTQ files for all BrainSeq Phase 1 subjects (n = 738) are available on Synapse [11, 49]. Genotypes are available from Globus [11] with restricted access. More information on the BrainSeq publicly available data can be found at http://eqtl.brainseq.org/.

Project name: dRFEtools

Project home page: https://github.com/paquolalab/dRFEtools

Archived version: https://doi.org/10.5281/zenodo.6884185 (v0.2.0)

Operating system(s): Linux, Mac OS, Windows

Programming language: Python

Other requirements: Python 3.7 or higher

License: GPL-3.0

### Competing interests

The authors declare no competing interests.

### Funding

This work is supported by the Lieber Institute for Brain Development and the National Institute on Minority Health and Health Disparities of the National Institutes of Health (K99MD016964) to K.J.M.B.

### Authors’ contributions

Conceptualization, K.J.M.B. and A.C.M.P; Methodology, K.J.M.B. and A.C.M.P.; Software, K.J.M.B., T.K., and A.C.M.P.; Formal Analysis, K.J.M.B; Writing – Original Draft, K.J.M.B.; Writing – Review & Editing, K.J.M.B. and A.C.M.P.; Visualization, K.J.M.B.; Supervision and Project Administration, A.C.M.P.; Funding Acquisition, K.J.M.B. and A.C.M.P.

## Acknowledgements

We would like to extend our deepest appreciation to our colleagues in the Erwin laboratory group for their comments and suggestions in the development of this software.

## Supplementary Information

### Data

#### Data S1: optimization.ipynb

Jupyter notebook with example code for performing optimization with random forest classification and regression for binary, multiclass, and regression analysis using simulated data.

#### Data S2: dRFEtools_binary_classification_results.tar.gz

Compressed tar zipped directory containing results from binary classification of major depression disorder or schizophrenia compared with neurotypical controls with dRFEtools using random forest classification. These results include text files with predictive features (minimum and redundant subsets), summary results across folds, and individual fold level results. Individual level results includes dRFEtools training and test scores, prediction results by individual, and full ranking list.

#### Data S3: dRFEtools_mdd_classification_GO_analysis.xlsx

Excel file containing gene term enrichment results for binary classification of major depression disorder using GOATools.

#### Data S4: dRFEtools_multiclassification_results.tar.gz

Compressed tar zipped directory containing results from multi -classification of neuropsychiatric disorders (major depression disorder, schizophrenia, or neurotypical controls) with dRFEtools using random forest classification. These results include text files with predictive features (minimum and redundant subsets), summary results across folds, and individual fold level results. Individual level results includes dRFEtools training and test scores, prediction results by individual, and full ranking list.

#### Data S5: dRFEtools_gene_imputation_results.tar.gz

Compressed tar zipped directory containing results from gene expression imputation of 69 non-mitochondria genes predictive in the multi-class classification. This includes compressed text files of dRFE individual level feature elimination results, dRFEtools training and test scores, full ranking list, and test predictions per feature. In addition to this, summary files and predictive features results (minimum and redundant subsets) are also contained in this compressed directory.

---

**Figure S1:**
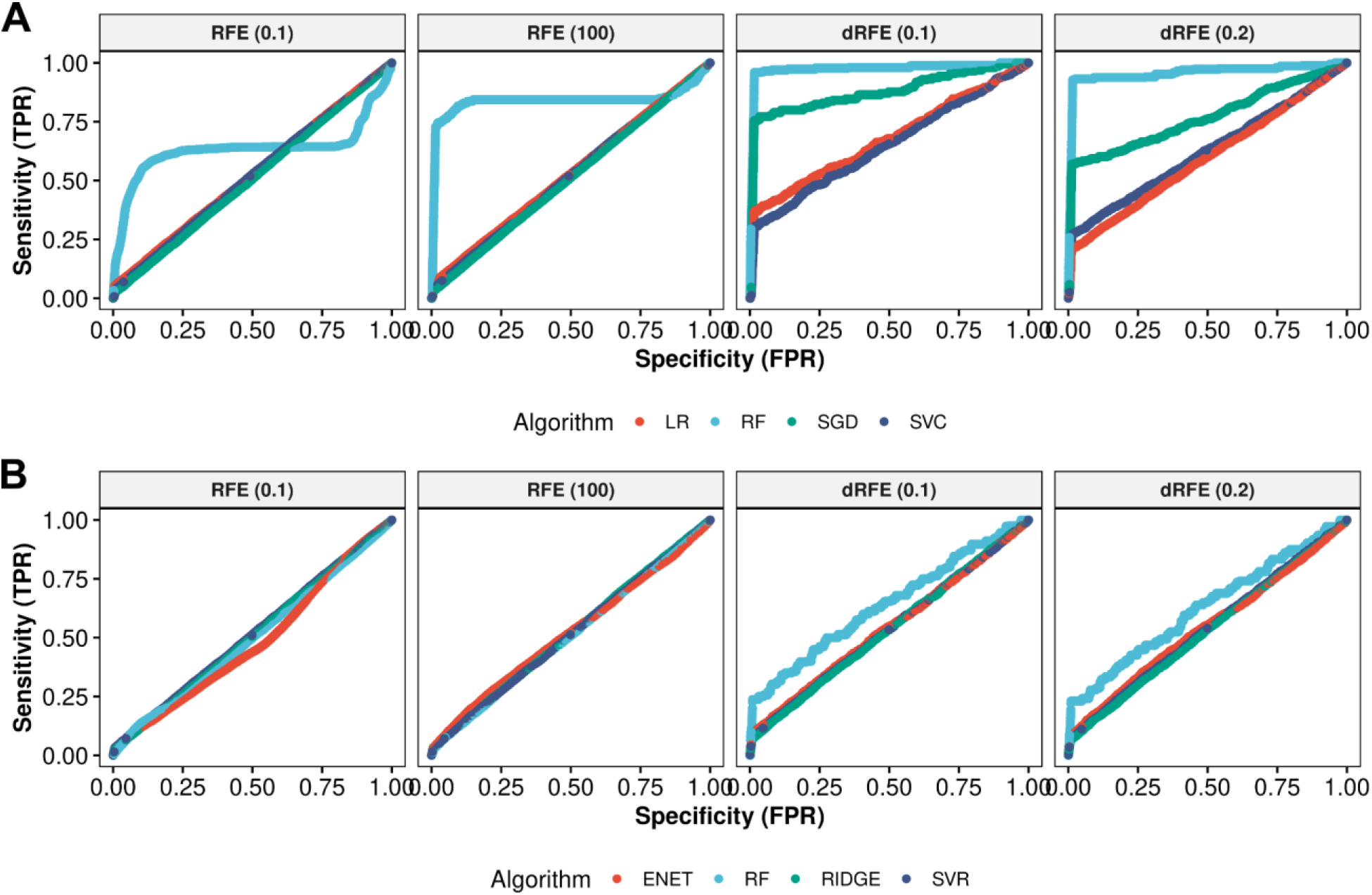
dRFEtools has improved feature selection sensitivity and specificity compared with current RFE algorithms using scikit-learn simulated data. ROC curve showing median true positive and false positive rate across 10-fold cross-validated simulated **A.** classification and **B.** regression data. LR: logistic regression, RF: random forest, SGD: stochastic gradient descent, ENET: elastic net, SVC: support vector classifier, and SVR: support vector regressor. RFE (0.1): RFE with 10% representing a static number equal to 10% of total features (2000 features each iteration). RFE (100): RFE that eliminates 100 features each iteration.

**Figure S2:**
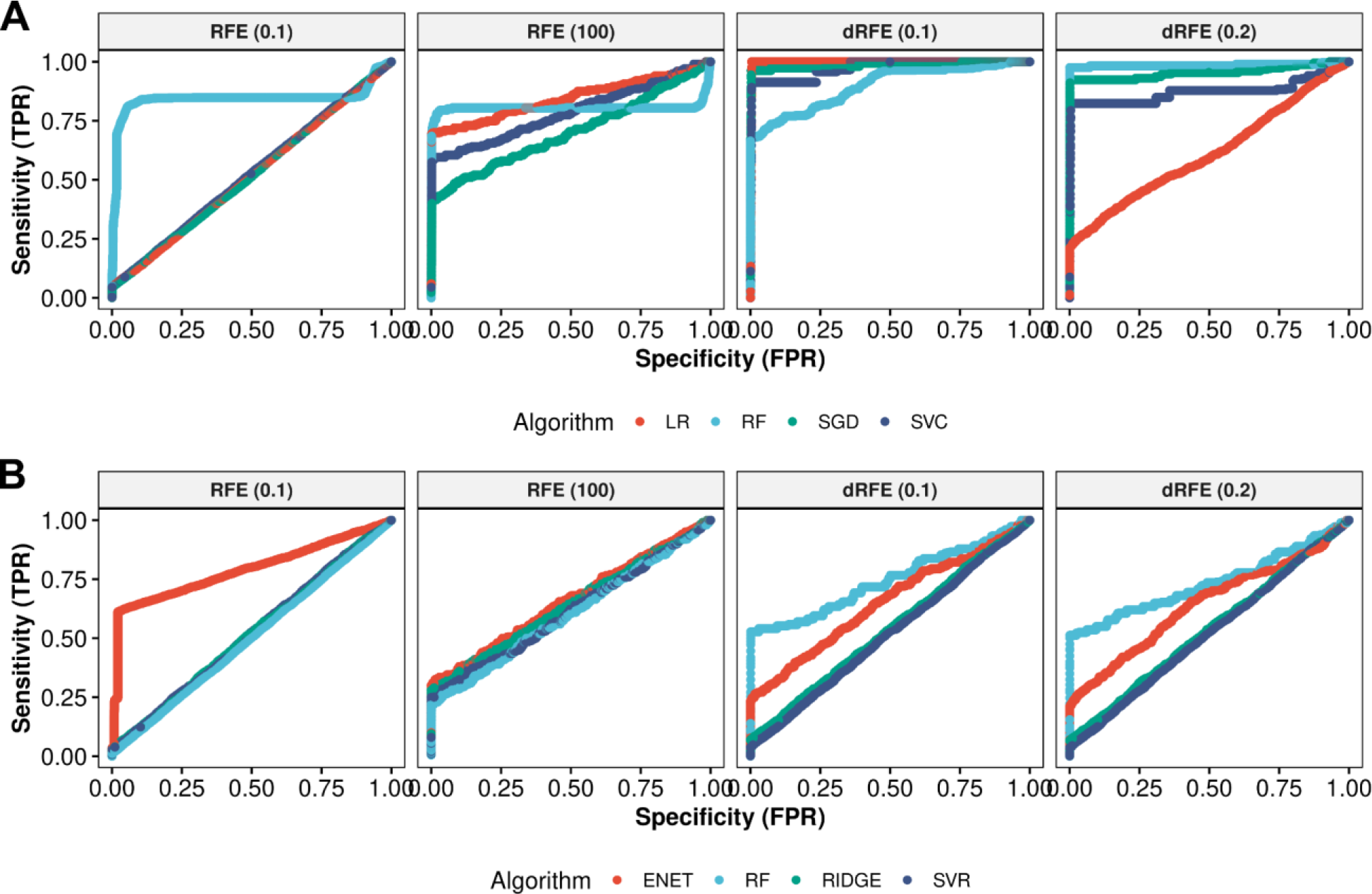
dRFEtools generally improves feature selection sensitivity and specificity compared with current RFE algorithms using biological simulated data. ROC curve showing median true positive and false positive rate across 10-fold cross-validated simulated **A.** bulk RNA-sequencing classification and **B.** quantitative trait regression data. LR: logistic regression, RF: random forest, SGD: stochastic gradient descent, ENET: elastic net, SVC: support vector classifier, and SVR: support vector regressor. RFE (0.1): RFE with 10% representing a static number equal to 10% of total features (2000 features each iteration). RFE (100): RFE that eliminates 100 features each iteration.

**Figure S3:**
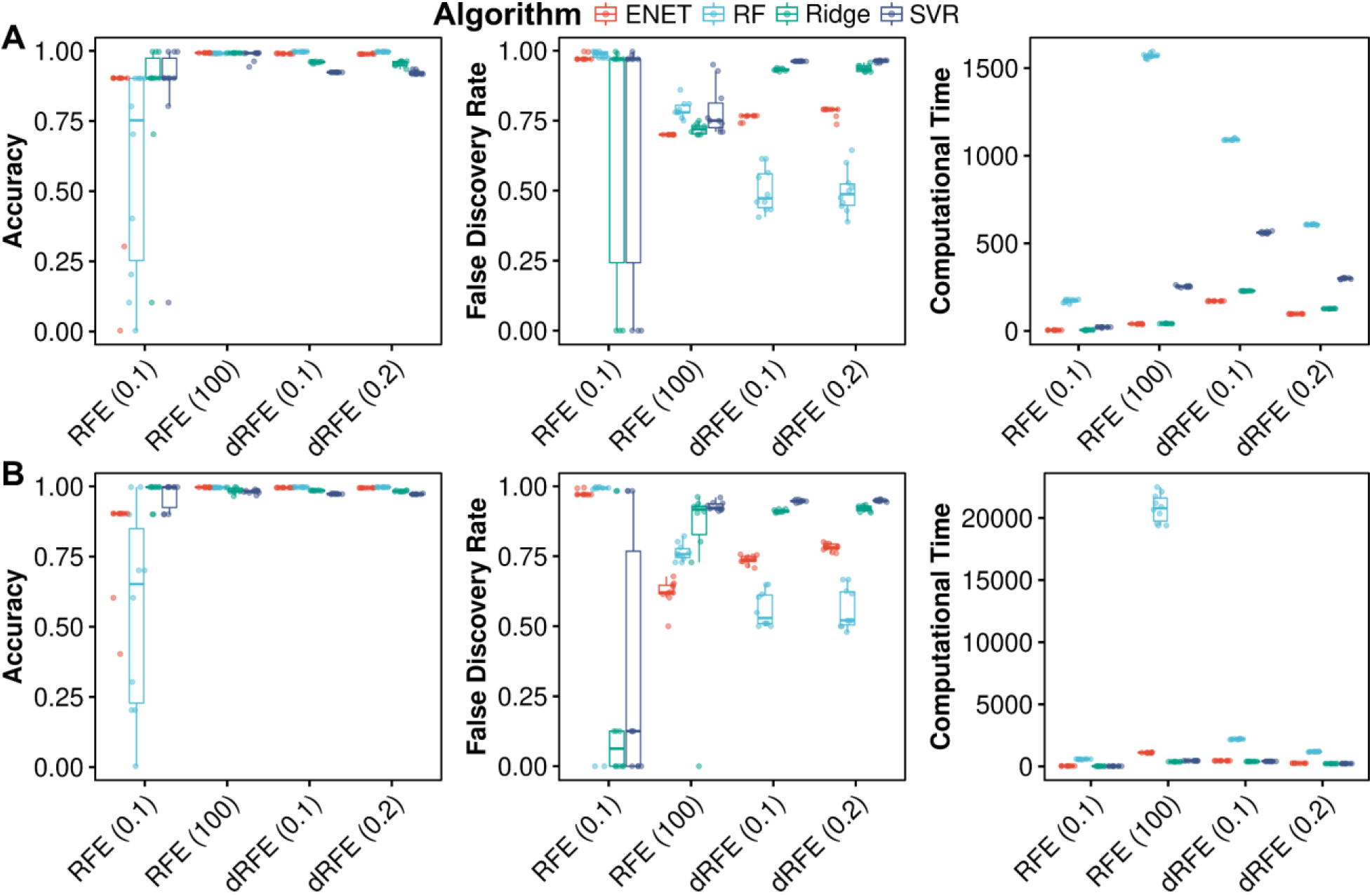
Comparison dRFEtools and RFE with and without one-hot encoding. Regression simulation with 10-fold cross-validation showing impact on computational time **A.** without and **B.** with one-hot encoding. LR: logistic regression, RF: random forest, SGD: stochastic gradient descent, ENET: elastic net, SVC: support vector classifier, and SVR: support vector regressor. Accuracy: feature selection accuracy. FDR: feature selection false discovery rate.

**Figure S4:**
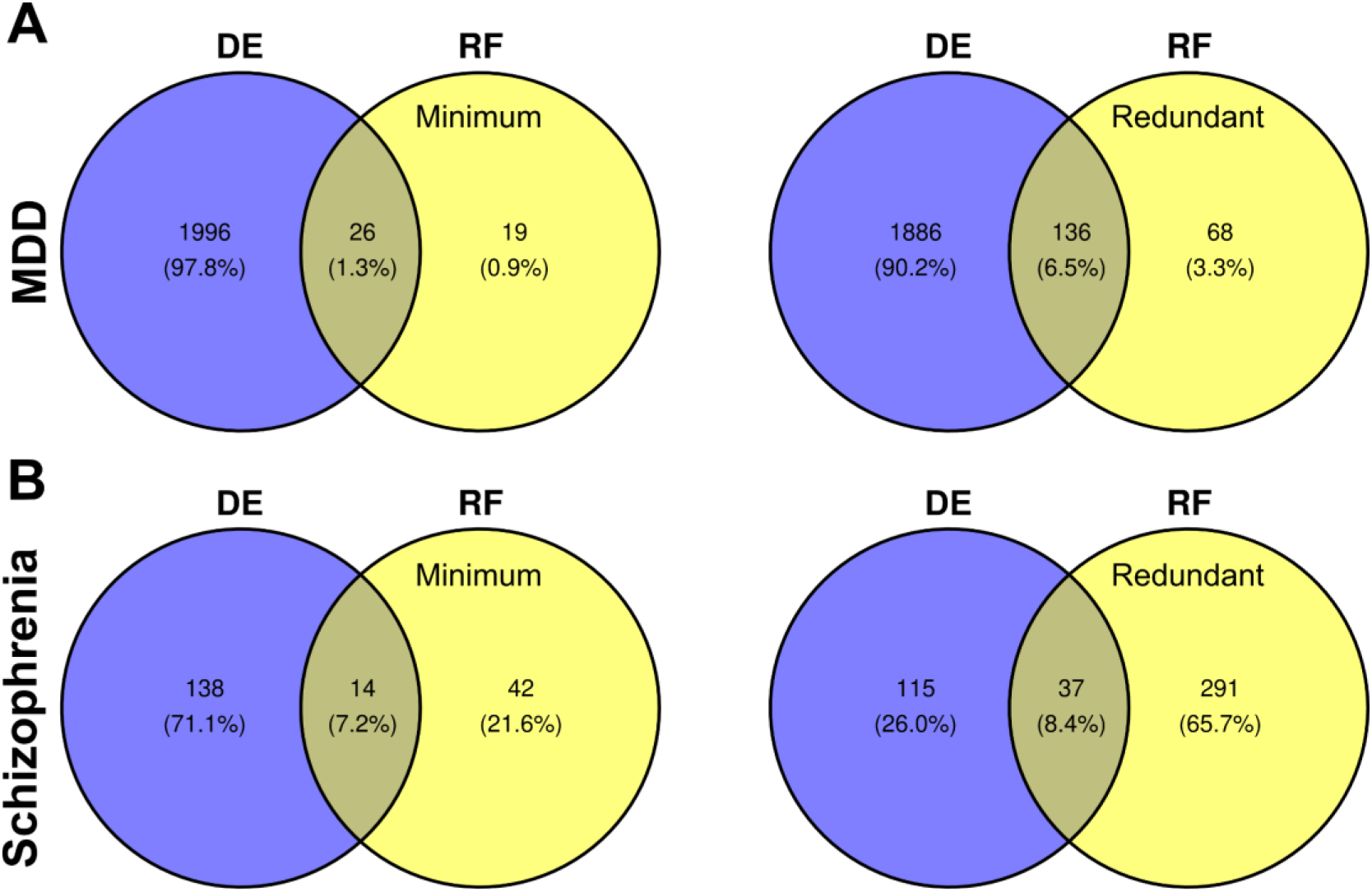
Higher overlap between traditional differential expression analysis and dRFE applied random forest classification associated with higher prediction accuracy. Venn diagrams showing overlap between DE (differential expression) using limma-voom and dRFEtools feature selection with random forest binary classification (RF) of the smallest (minimum) predictive set (left) and the redundant predictive set (right) for **A.** major depression disorder (MDD) and **B.** schizophrenia.

**Figure S5:**
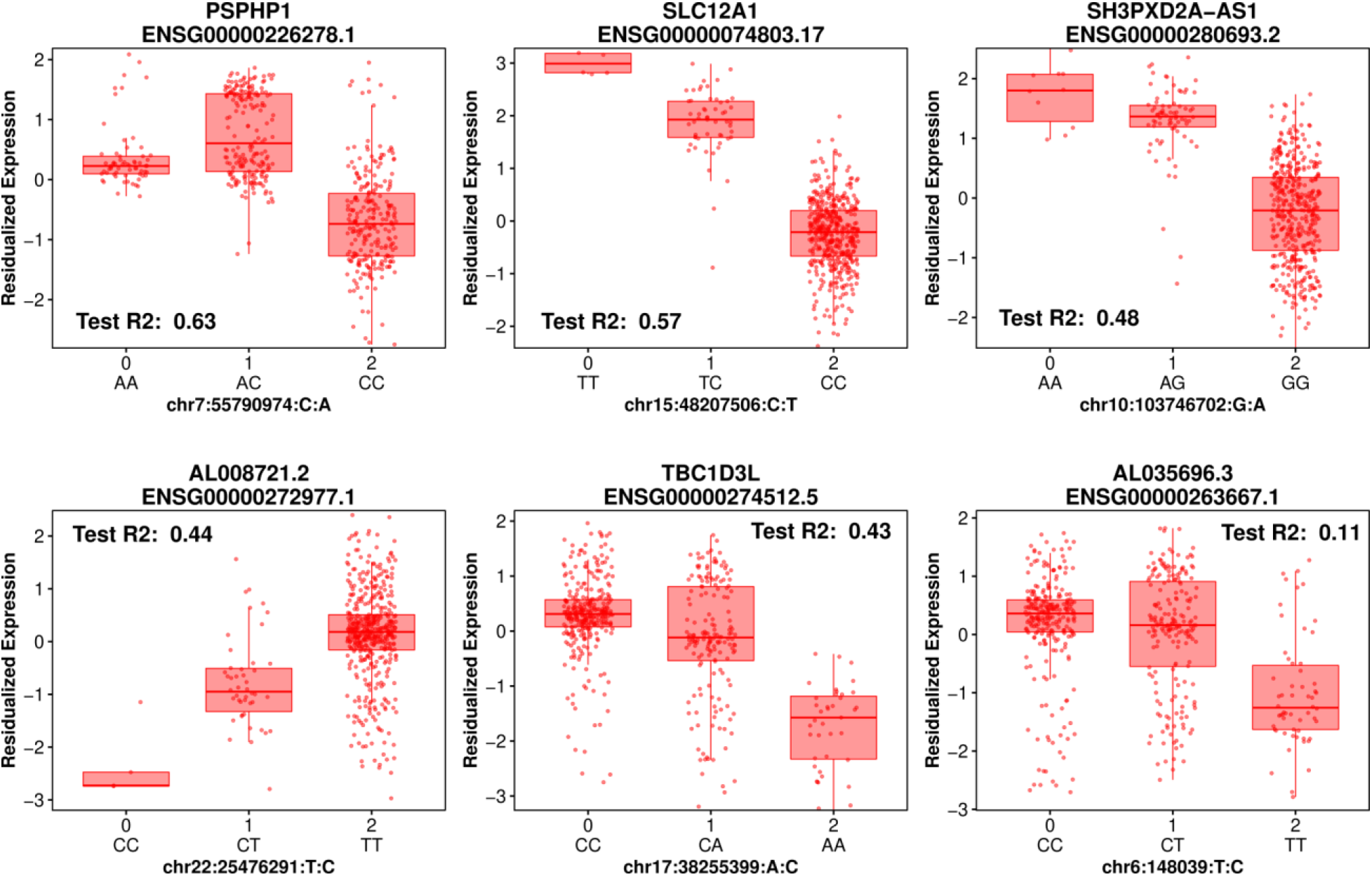
dRFEtools selects features with strong genetic association to gene expression similar to expression quantitative trait loci (eQTL) analysis. Boxplots showing all significant (random forest regression, test r^2^ score > 0) association between genetic variants and gene expression. Median test r^2^ score across 10-fold cross-validation annotated in plots.

**Figure S6:**
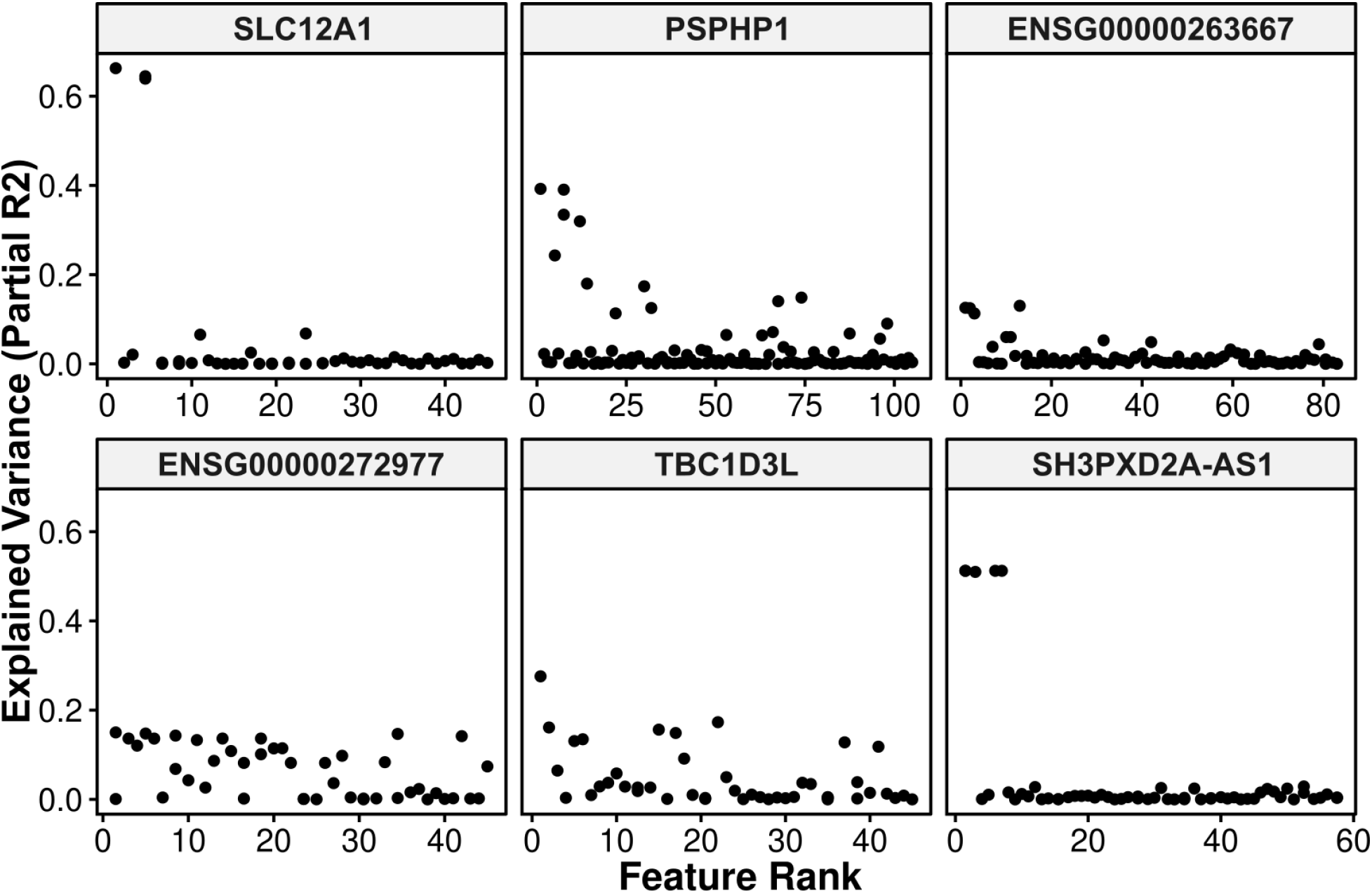
Feature rank associates with explained variance. Scatterplot showing median feature rank and explained variance.

## Tables

**Table S1.** A sample breakdown for adults (age > 17), BrainSeq (Phase 1) postmortem DLPFC [11] by diagnosis. Abbreviations: Female (F), Male (M), African Ancestry (AA), European Ancestry (EA), and RNA integrity number (RIN).

| Diagnosis | Sample Size | Sex | Race | Age<br>(mean ± sd) | RIN<br>(mean ± sd) |
| --- | --- | --- | --- | --- | --- |
| Neurotypical control | 207 | 60F/147M | 117AA/90EA | 43.4 ± 15.4 | 8.3 ± 0.7 |
| Major depression disorder | 142 | 56F/86M | 15AA/127EA | 44.8 ± 13.2 | 8.0 ± 0.9 |
| Schizophrenia | 172 | 62F/110M | 77AA/95EA | 49.7 ± 14.8 | 8.0 ± 0.9 |

## Notes

https://pypi.org/project/drfetools/

### Competing Interest Statement

The authors have declared no competing interest.

