## Supplementary Data 1: Optimization jupyter notebook for "dRFEtools: Dynamic recursive feature elimination for omics": optimization.ipynb

```
# %% [markdown]
# Example of optimizing lowess fit and max and redundant set selection

# %% [code]
import os,errno
import dRFEtools
import numpy as np
import pandas as pd
from sklearn.model_selection import KFold
from sklearn.datasets import make_regression
from sklearn.datasets import make_classification
from sklearn.model_selection import StratifiedKFold
from sklearn.model_selection import train_test_split

# %% [code]
def mkdir_p(directory):
    try:
        os.makedirs(directory)
    except OSError as e:
        if e.errno != errno.EEXIST:
            raise

# %% [markdown]
## Optimize classification

# %% [code]
outdir = 'classification/'
mkdir_p(outdir)

# %% [code]
# Create a dataset with only 10 informative features
X, y = make_classification(
    n_samples=500, n_features=20000, n_informative=100, n_redundant=300,
    n_repeated=0, n_classes=2, n_clusters_per_class=1, random_state=13
)

cv = StratifiedKFold(n_splits=5, shuffle=True, random_state=13)
#cla = dRFEtools.LogisticRegression(max_iter=1000, n_jobs=-1)
cla = dRFEtools.RandomForestClassifier(n_estimators=100, n_jobs=-1, 
                                      oob_score=True, random_state=13)

# %% [code]
fold = 1
for train_index, test_index in cv.split(X, y):
    X_train, X_test = X[train_index], X[test_index]
    y_train, y_test = y[train_index], y[test_index]
    fold += 1
fold -= 1

features = ["feature_%d" % x for x in range(X_train.shape[1])]
d, pfirst = dRFEtools.rf_rfe(cla, X_train, y_train, np.array(features), 
                            fold, outdir, elimination_rate=0.1, RANK=False)

# %% [markdown]
### Optimize lowess fit: fraction of data to use when estimating y-values

# %% [code]
## Single Fold examination
for frac in [0.1, 0.15, 0.2, 0.25, 0.3, 0.35, 0.4]:
    dRFEtools.optimize_lowess_plot(d, fold, outdir, frac=frac, step_size=0.02, 
                                   classify=True, save_plot=True)

# %% [markdown]
### Optimize redundant selection: examine rate change

# %% [code]
for step_size in [0.01, 0.02, 0.03, 0.04, 0.05, 0.10]:
    dRFEtools.optimize_lowess_plot(d, fold, outdir, frac=0.3, step_size=step_size, 
                                   classify=True, save_plot=True)

# %% [code]
## Highest value (without smoothing)
max(d, key=lambda x: d[x][1])

# %% [code]
print(dRFEtools.extract_max_lowess(d, frac=0.30))
## Using a conservative step size
dRFEtools.extract_redundant_lowess(d, frac=0.30, step_size=0.02)

# %% [markdown]
## Optimize regression

# %% [code]
outdir = 'regression/'
mkdir_p(outdir)

# %% [code]
# Create a dataset with only 10 informative features
X, y = make_regression(
    n_samples=500, n_features=20000, n_informative=400, bias=0.02,
    n_targets=1, noise=5, random_state=13
)
cv = KFold(n_splits=5, shuffle=True, random_state=13)
regr = dRFEtools.RandomForestRegressor(n_estimators=100, oob_score=True, 
                                       n_jobs=-1, random_state=13)

# %% [code]
fold = 1
for train_index, test_index in cv.split(X, y):
    X_train, X_test = X[train_index], X[test_index]
    y_train, y_test = y[train_index], y[test_index]
    fold += 1
fold -= 1

features = ["feature_%d" % x for x in range(X_train.shape[1])]
d, pfirst = dRFEtools.rf_rfe(regr, X_train, y_train, np.array(features), 
                            fold, outdir, elimination_rate=0.1, RANK=False)

# %% [markdown]
### Optimize lowess fit: fraction of data to use when estimating y-values

# %% [code]
## Single Fold examination
for frac in [0.1, 0.15, 0.2, 0.25, 0.3, 0.35, 0.4]:
    dRFEtools.optimize_lowess_plot(d, fold, outdir, frac=frac, step_size=0.02, 
                                   classify=False, save_plot=True)

# %% [markdown]
### Optimize redundant selection: examine rate change

# %% [code]
for step_size in [0.01, 0.02, 0.03, 0.04, 0.05, 0.1]:
    dRFEtools.optimize_lowess_plot(d, fold, outdir, frac=0.25, step_size=step_size, 
                                   classify=False, save_plot=True)

# %% [code]
## Highest values without smoothing
max(d, key=lambda x: d[x][1])

# %% [code]
print(dRFEtools.extract_max_lowess(d, frac=0.25))
dRFEtools.extract_redundant_lowess(d, frac=0.25, step_size=0.02)

# %% [markdown]
## Optimize classification: multi-class

# %% [code]
outdir = 'multiclass/'
mkdir_p(outdir)

# %% [code]
# Create a dataset with only 10 informative features
X, y = make_classification(
    n_samples=500, n_features=20000, n_informative=100, n_redundant=300,
    n_repeated=0, n_classes=4, n_clusters_per_class=1, random_state=13
)

cv = StratifiedKFold(n_splits=5, shuffle=True, random_state=13)
cla = dRFEtools.RandomForestClassifier(n_estimators=100, n_jobs=-1, 
                                      oob_score=True, random_state=13)

# %% [code]
fold = 1
for train_index, test_index in cv.split(X, y):
    X_train, X_test = X[train_index], X[test_index]
    y_train, y_test = y[train_index], y[test_index]
    fold += 1
fold -= 1

features = ["feature_%d" % x for x in range(X_train.shape[1])]
d, pfirst = dRFEtools.rf_rfe(cla, X_train, y_train, np.array(features), 
                            fold, outdir, elimination_rate=0.1, RANK=False)

# %% [markdown]
### Optimize lowess fit: fraction of data to use when estimating y-values

# %% [code]
## Single Fold examination
for frac in [0.1, 0.15, 0.2, 0.25, 0.3, 0.35, 0.4]:
    dRFEtools.optimize_lowess_plot(d, fold, outdir, frac=frac, step_size=0.02, 
                                   classify=True, multi=True, save_plot=True)

# %% [markdown]
### Optimize redundant selection: examine rate change

# %% [code]
for step_size in [0.01, 0.02, 0.03, 0.04, 0.05, 0.1]:
    dRFEtools.optimize_lowess_plot(d, fold, outdir, frac=0.25, step_size=step_size, 
                                   classify=True, multi=True, save_plot=True)

# %% [code]
## Highest value (without smoothing)
max(d, key=lambda x: d[x][1])

# %% [code]
print(dRFEtools.extract_max_lowess(d, frac=0.25))
dRFEtools.extract_redundant_lowess(d, frac=0.25, step_size=0.015)

# %% [code]
```
